# An anterior–posterior gradient in hippocampal subfield volumes linking sleep health to cognition in young adults

**DOI:** 10.1101/2025.07.30.666702

**Authors:** Tianfang Han, Tongfang Ding, Shubing Li, Toru Ishihara

## Abstract

Sleep plays a critical role in maintaining hippocampal integrity and supporting cognitive function. However, it remains unclear how variations in sleep are associated with the structurally heterogeneous subfields of the hippocampus and how this association relates to cognition in healthy young adults. This study aimed to elucidate the multivariate relationships among sleep duration, quality, and timing, the specific hippocampal subfield volumes, and cognitive functions in a large sample of healthy young adults. We applied multiset canonical correlation analysis to data from 942 young adults (ages 22–37) from the Human Connectome Project. We examined relationships among self-reported sleep parameters (via the Pittsburgh Sleep Quality Index), high-resolution hippocampal subfield volumes, and a broad set of cognitive performance measures. We identified a single significant mode of covariation, wherein better sleep health—characterized by a moderate sleep duration (reflecting a nonlinear, inverted U-shaped relationship), higher sleep quality (e.g., fewer awakenings, less discomfort), and optimal sleep timing—was robustly associated with larger hippocampal volumes, particularly in the anterior subfields. This sleep–hippocampus pattern was also associated with enhanced cognitive performance. Our findings reveal a specific link among sleep, an anterior-dominant pattern of hippocampal structure, and cognition in healthy young adulthood. The anterior hippocampus may serve as a key neural hub through which the cognitive benefits of healthy sleep are manifested early in life.

## Introduction

The hippocampus is a critical brain structure involved in the encoding and consolidation of new memories (Battaglia et al., 2011; Shapiro & Eichenbaum, 1999; Sridhar et al., 2023). Its role extends beyond memory formation, underpinning diverse domains of human cognition such as flexible information processing, spatial navigation, social cognition, and executive functioning (Maguire et al., 2000; Montagrin et al., 2018; O’Shea et al., 2016; Rubin et al., 2014; Trinkler et al., 2009). The human hippocampus shows remarkable neuroplasticity, undergoing structural and functional changes in response to various lifestyle factors (Binnewies et al., 2023; Fotuhi et al., 2012; Namsrai et al., 2023; Popa-Wagner et al., 2020). Among these factors, sleep appears to play a foundational role in supporting cognitive health (Paller et al., 2021). Chronic sleep insufficiency has been associated with accelerated hippocampal volume loss, especially in individuals reporting poor sleep quality and greater daytime fatigue (Fjell et al., 2020). A recent meta-analysis further showed that inadequate sleep duration and low sleep quality are associated with smaller bilateral hippocampal volumes (Namsrai et al., 2025).

However, the relationship between sleep and the hippocampus is likely more nuanced at the subfield level. The hippocampus is a heterogeneous structure composed of subfields—including the cornu ammonis (CA), dentate gyrus (DG), and subiculum—each characterized by distinct cytoarchitecture, connectivity, and functional roles (Bast, 2007; Dekraker et al., 2021; Genon et al., 2021). These subfields are organized along multiple anatomical axes, most notably the anterior– posterior and medial-lateral axes, each with distinct functional specializations (Plachti et al., 2019; Strange et al., 2014). Thus, understanding how sleep differentially affects individual subfields is critical for uncovering the mechanisms of hippocampal vulnerability and resilience.

Nevertheless, studies examining the link between sleep and hippocampal subfield volumes have produced inconsistent findings. Early research in clinical populations, such as those with post-traumatic stress disorder or insomnia, indicated that poor sleep was associated with volume reductions in subfields like CA1, CA3, and DG, although the specific regions affected varied by condition and symptom profile (Joo et al., 2014; Neylan et al., 2010). Subsequent studies in non-clinical aging and developmental samples further complicated the picture. Some reported that poor sleep quality or atypical sleep duration (both short and long) were linked to atrophy in a broad range of subfields, including the CA regions, subiculum, and the DG (De Looze et al., 2022; Liu et al., 2021). Others, in contrast, found no significant associations in certain subfields (e.g., CA1, DG), or even linked longer sleep duration to smaller volumes in areas such as the presubiculum (Mourtzi et al., 2024).

Several factors may account for these inconsistencies. Methodological differences—including variations in how sleep is assessed (e.g., quantity vs. quality), subfield segmentation techniques, and small sample sizes—likely contribute to the variability (Kim et al., 2016; Teicher et al., 2017). Additionally, the presence of confounding factors, such as underlying clinical conditions, complicates efforts to isolate the unique effects of sleep (Zhou et al., 2025). Importantly, most previous studies have focused exclusively on anatomical changes, leaving a critical gap in understanding how sleep-related differences in hippocampal subfields are linked to cognitive performance.

To address these gaps, we used a large and well-characterized dataset from the Human Connectome Project (HCP) (Van Essen et al., 2013). Using a multivariate analytical approach, we aimed to simultaneously examine the complex relationships among sleep parameters (duration, quality, and timing), high-resolution hippocampal subfield volumes, and diverse cognitive functions in healthy young adults. We hypothesized that better sleep would be associated with larger hippocampal volumes and enhanced cognitive performance. Given the exploratory nature of subfield-specific associations, we also sought to identify which aspects of sleep are most strongly related to particular subfields, and how these anatomical patterns covary with cognition. Further, we examined whether these patterns were organized along the two primary functional gradients of the hippocampus: the anterior-posterior and medial-lateral axes (Plachti et al., 2019; Strange et al., 2014).

## Methods

### Participants

We used data from the HCP, which initially included 1,206 individuals aged 22–37 years. Initiated in 2010, the HCP aims to provide a comprehensive understanding of human brain connectivity and function, allowing integrated comparisons of neural circuits, behaviors, and genetic profiles (Van Essen et al., 2013). Behavioral and 3T magnetic resonance imaging (MRI) data used in this study were collected between 2012 and 2015. Exclusion criteria included a history of severe psychiatric illness or head trauma, evidence of neurological or cardiopulmonary disease, pregnancy, or elevated heavy metal exposure. The study protocol was approved by the Institutional Review Board of Washington University in St. Louis (Office for Human Research Protections, No. 201204036), and all procedures were conducted in accordance with the Declaration of Helsinki. Written informed consent was obtained from all participants. From the initial sample of 1,206 participants, we excluded individuals based on the HCP quality control guidelines, including those flagged for anatomical anomalies (Issue code A), segmentation and surface errors (Issue code B), or data acquisition issues (Issue code C). We also excluded participants with missing data for the variables of interest (sleep, cognitive, or MRI data) using listwise deletion. This resulted in a final sample of 942 participants for the main analyses.

### Sleep assessment

Sleep quality and disturbances were assessed using the Pittsburgh Sleep Quality Index (PSQI) (Buysse et al., 1989). The PSQI is a validated self-report questionnaire that evaluates sleep quality and disturbances over the past month, consisting of seven component scores derived from 19 items. The component scores are summed to generate a global score ranging from 0 to 21, with higher scores indicating poorer sleep quality. To enable a more granular analysis of the associations between sleep and hippocampal structure while minimizing redundancy, we focused on individual PSQI items. For sleep duration, given prior reports of a quadratic association with hippocampal volume (Tai et al., 2022), both the centered mean and its squared term were included to model a potential nonlinear effect. To accommodate the circular nature of sleep timing variables, self-reported bedtimes and wake times were transformed into two-dimensional unit vectors using sine and cosine functions. Each time point was first converted to a radian value on the 24-hour clock, then mapped to the unit circle via cos(θ) and sin(θ). These transformed variables were entered as continuous predictors, preserving temporal directionality for linear multivariate modeling.

### MRI acquisition

All neuroimaging data were acquired using a customized 3T Siemens Skyra scanner dedicated to the HCP, equipped with a 32-channel head coil. High-resolution T1-weighted (T1w) and T2-weighted (T2w) structural images were obtained following standardized HCP acquisition protocols (Glasser et al., 2013). T1w images were collected using a 3D magnetization-prepared rapid acquisition gradient echo (MPRAGE) sequence (Mugler & Brookeman, 1990) with two averages and the following parameters: 0.7 mm isotropic voxel resolution (FOV = 224 mm; matrix size = 320; 256 sagittal slices), repetition time (TR) = 2400 ms, echo time (TE) = 2.14 ms, inversion time (TI) = 1000 ms, flip angle = 8°, bandwidth = 210 Hz/pixel, and echo spacing = 7.6 ms. T2w images were collected using a variable flip angle turbo spin-echo sequence (Siemens SPACE) (Mugler et al., 2000), matched to the T1w geometry, with TR = 3200 ms, TE = 565 ms, and bandwidth = 744 Hz/pixel. Two averages were obtained for each image type.

### MRI data preprocessing

We used FreeSurfer version 8.0.0 to process structural MRI data, employing the standard recon-all pipeline for cortical and subcortical segmentation (Fischl, 2012). To enhance the accuracy of the anatomical reconstruction, both T1w and T2w images were utilized as inputs. The inclusion of T2w data is particularly effective for improving the delineation of the pial surface and correcting topological defects, leading to more reliable cortical models. For subjects with multiple available scans, all T1w and T2w images were leveraged by recon-all to increase the signal-to-noise ratio and improve the robustness of the final segmentation. Following the recon-all processing, hippocampal subfield segmentation was conducted using the dedicated automated algorithm within FreeSurfer, which applies a probabilistic atlas-based approach informed by ultra-high resolution ex vivo MRI and histological data (Iglesias et al., 2015). This method allows for the parcellation of the hippocampus into anatomically defined subfields, including CA1, CA2/3, CA4, DG, subiculum, presubiculum, fimbria, and hippocampal-amygdaloid transition area (HATA), among others.

### Cognitive function assessment

Thirteen cognitive tests were administered to assess diverse cognitive functions, including episodic memory (Picture Sequence Memory), cognitive flexibility (Dimensional Change Card Sort), inhibitory control (Flanker task), fluid intelligence (Penn Progressive Matrices), language/reading decoding (Oral Reading Recognition), language/vocabulary comprehension (Picture Vocabulary), processing speed (Pattern Completion Processing Speed), self-regulation/impulsivity (Delay Discounting), spatial orientation (Variable Short Penn Line Orientation Test), sustained attention (Short Penn Continuous Performance Test), verbal episodic memory (Penn Word Memory Test), emotion recognition (Penn Emotion Recognition Test) and working memory (List Sorting). A detailed description of all cognitive measures and derived variables used in the analysis is provided in Supplementary Table S1. Composite scores for Fluid Cognition, Early Childhood, Crystallized Cognition, and overall Cognitive Function were calculated from these measures.

### Statistical analysis

All statistical analyses were conducted using R statistical software (version 4.5.0). To investigate multivariate relationships among three sets of variables—sleep parameters, hippocampal subfield volumes, and cognitive functions—we employed multiset canonical correlation analysis (MCCA). MCCA extends conventional canonical correlation analysis to identify associations across more than two data modalities simultaneously (Zhuang et al., 2020). MCCA was performed using modified versions of the MultiCCA functions from the PMA package. Prior to MCCA, dimensionality reduction was applied to each variable set using principal component analysis (PCA) via the prcomp function. For each of the three variable sets, we selected the minimum number of principal components required to explain at least 80% of the total variance. The statistical significance of the MCCA components was assessed using a non-parametric permutation test (n = 1,000). Age, sex, and handedness were included as nuisance covariates and regressed from the data before performing PCA. Further, to examine region-specific associations beyond overall brain size, we conducted additional analyses including intracranial volume (ICV) as a covariate. To formally test for the observed anatomical gradients, we examined whether the canonical variate loadings of the hippocampal subfields varied systematically along the anterior-posterior and medial-lateral axes. Using linear multiple regression with robust standard errors (lm_robust function from the estimatr package in R), we regressed the subfield loadings onto their Montreal Neurological Institute (MNI) space coordinates. The model included terms for the anterior-posterior axis (Y-coordinate), the medial-lateral axis (absolute X-coordinate), and hemisphere (left vs. right) to quantify their respective contributions to the observed pattern of associations. To assess the robustness of our results, we conducted sensitivity analyses by varying the number of principal components included and incorporating additional covariates, such as years of education, income, body mass index, systolic and diastolic blood pressure, physical fitness (sub-maximal cardiovascular endurance, gait speed, hand dexterity, and grip strength), and alcohol and tobacco use, during the regression step. Statistical tests were repeated across these model variations to confirm the stability and consistency of our findings.

## Results

### Participant characteristics

Table 1 provides a comprehensive overview of the demographic characteristics of the study cohort. The final sample consisted of 942 individuals, with 516 females (55%). The mean chronological age was 29 years. The mean amount of sleep and PSQI score were 6.8 hours and 4.8, respectively.

**Table 1.** Demographic and key behavioral characteristics of the study cohort.

| Variables | N (%) or mean (SD) | Range |
| --- | --- | --- |
| Gender |  |  |
| Female | 516 (55%) |  |
| Male | 426 (45%) |  |
| Age | 29 (4) | 22 to 37 |
| Handedness (Laterality Quotient) | 66 (44) | -100 to 100 |
| Right (Laterality Quotient > 40) | 801 (85%) |  |
| Left (Laterality Quotient < -40) | 57 (6%) |  |
| Mixed ($-40 \leq \text{Laterality Quotient} \leq +40$) | 84 (9%) | |
| Sleep<sup>a</sup> |  | 0-21 |
| Good sleep (PSQI_Score: 0-4) | 506 (54%) | 0 to 4 |
| Mild sleep disturbance (PSQI_Score: 5-10) | 393 (42%) | 5 to 10 |
| Moderate sleep disturbance (PSQI_Score: 11-15) | 41 (4%) | 11 to 15 |
| Severe sleep disturbance (PSQI_Score: 16-21) | 2 (0%) | 16 to 21 |
| Cognitive task performance<sup>b, c</sup> |  |  |
| Fluid Cognition Composite score | 115 (12) | 85 to 145 |
| Early Childhood Composite score | 117 (11) | 86 to 154 |
| Cognitive Function Composite score | 112 (15) | 85 to 153 |
| Crystallized Cognition Composite score | 118 (10) | 90 to 154 |

Note: Percentages may not total 100 because of rounding. <sup>a</sup> PSQI Global Score and sleep quality categories are reported. <sup>b</sup> The raw score is standardized based on the age-appropriate band of the Toolbox Norming Sample (age bands: 18-29, 30-35), with a score of 100 representing the national average performance. A score of 115 or 85 indicates performance one standard deviation above or below the national average within each age band. To understand the fitness level of the target population relative to the national population, we used age-standardized scores here, while unstandardized values were used for statistical analyses. <sup>c</sup> Only composite scores are reported.

### Main analysis

The MCCA revealed a single significant mode that captured the shared variance among sleep parameters, hippocampal subfield volumes, and cognitive functions (Figure 1A; sum of canonical correlation coefficients = 0.80; *p* < 0.001, permutation test). The correlations between the original variables and their respective canonical variates are presented in Figures 1B, 1E, and 2B.

**Figure 1.**
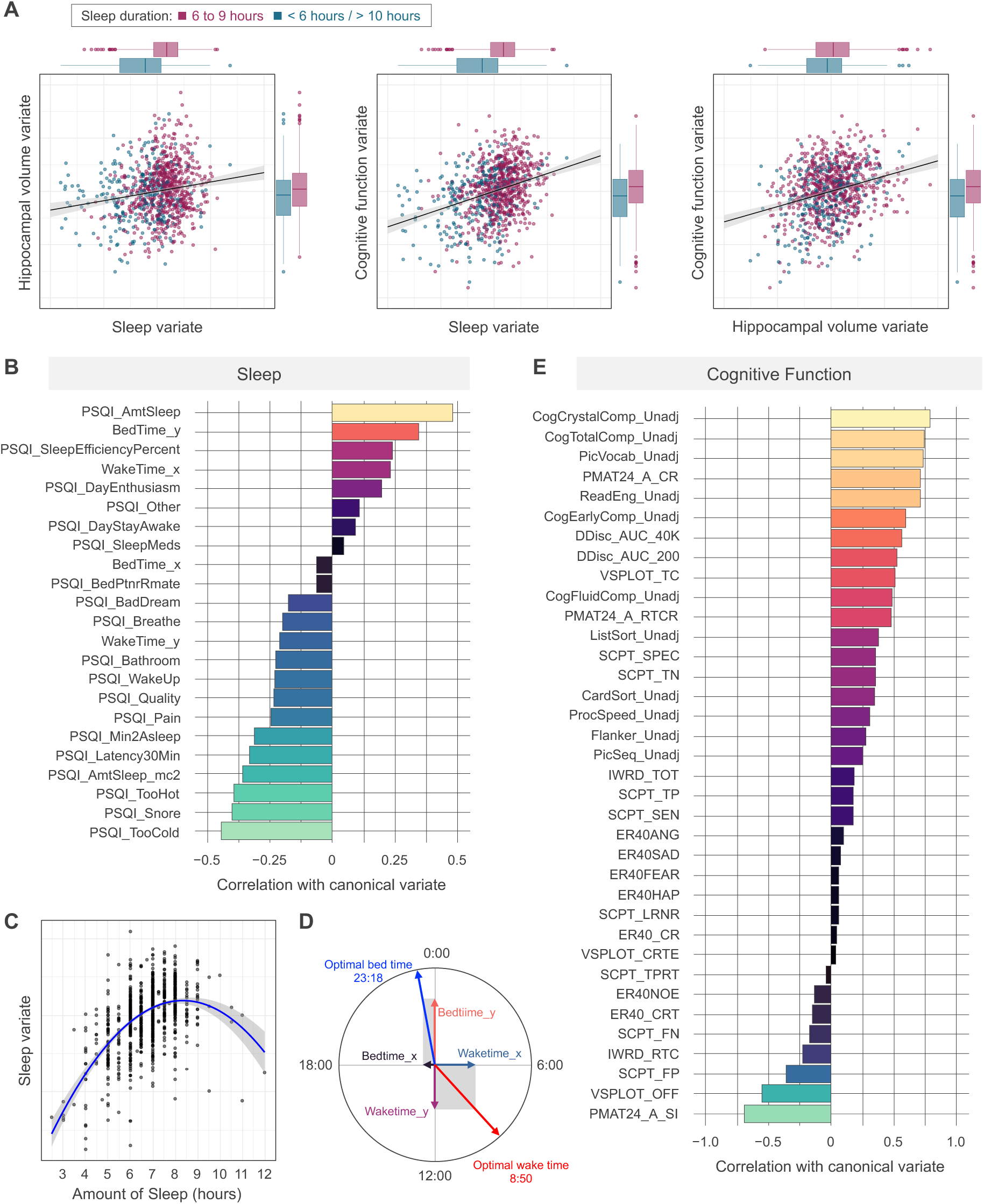
Multiset canonical correlation analysis of sleep, hippocampal volume, and cognitive function. (A) Scatter plots illustrating the pairwise relationships between the canonical variates for sleep, hippocampal volume, and cognitive function. Colors indicate sleep duration groups. (B) Correlations of individual sleep parameters with the sleep canonical variate. Positive values indicate an association with a profile of healthier sleep (e.g., higher efficiency, moderate duration). (C) A nonlinear, inverted U-shaped relationship between sleep duration and the sleep canonical variate. The blue line represents the quadratic fit, with the gray shading indicating the 95% confidence interval. (D) A circular plot visualizing optimal sleep timing. The red arrow indicates the optimal time vector, corresponding to a bedtime of approximately 23:18 and a wake time of approximately 08:50. (E) Correlations of individual cognitive test scores with the cognitive function canonical variate. Positive values indicate an association with better cognitive performance.

**Figure 2.**
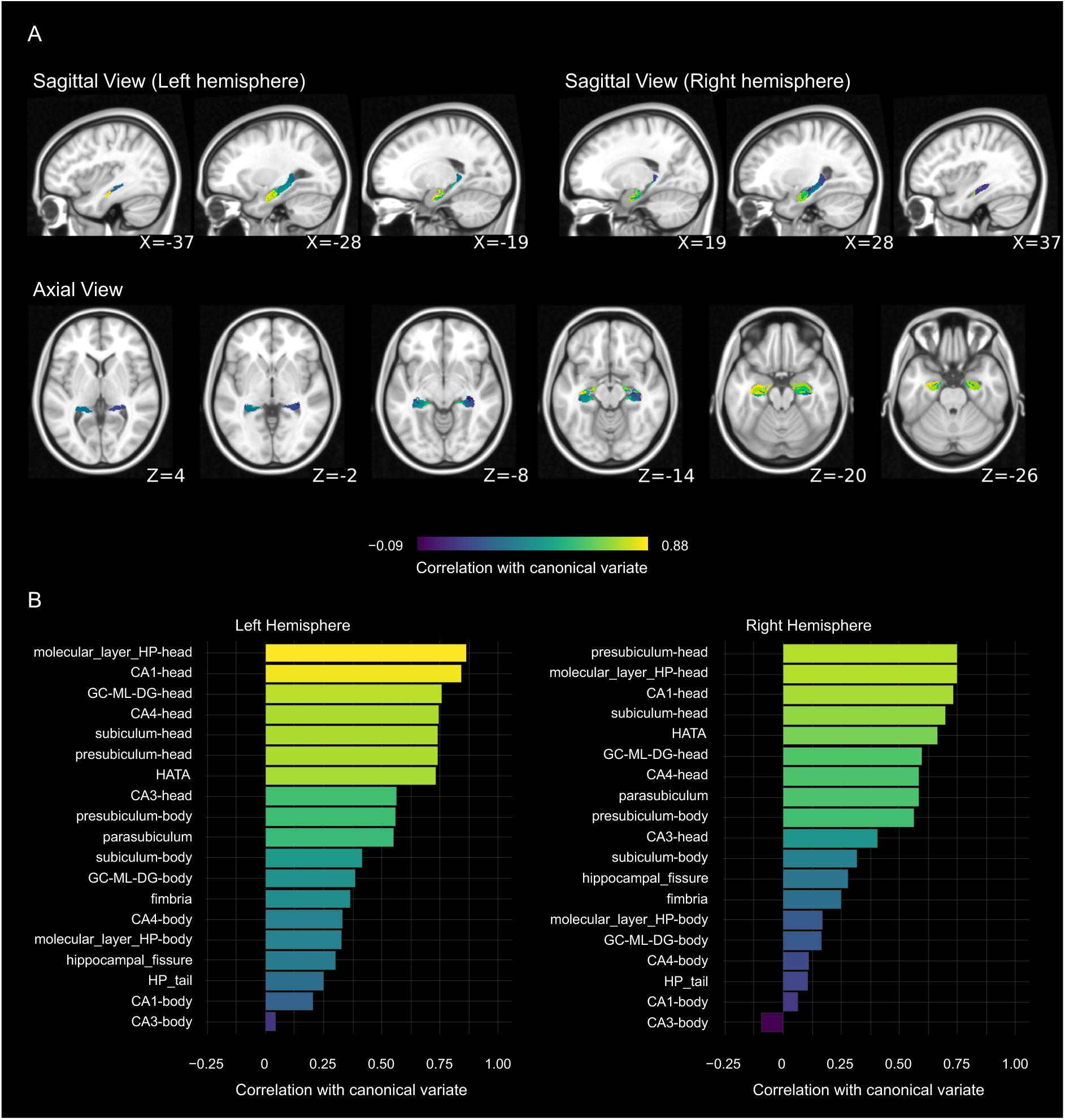
Spatial distribution of correlations between hippocampal subfield volumes and the canonical variate. (A) Sagittal and axial views of a representative hippocampus showing the spatial distribution of correlations between subfield volumes and the hippocampal canonical variate. The color bar indicates the strength of the correlation. (B) Bar plots showing the correlation coefficients for individual hippocampal subfields in the left and right hemispheres with the hippocampal canonical variate.

For sleep parameters, total sleep duration showed a strong positive linear association with the canonical variate (r = 0.48), while its squared term showed a negative association (r = –0.36), collectively indicating a nonlinear, inverted U-shaped relationship where both short and long sleep durations were linked to a less optimal profile (Figure 1C). Sleep efficiency showed a positive association with the canonical variate (r = 0.24). In addition to sleep quantity, several indicators of poor sleep quality were negatively associated with the canonical variate, including snoring (r = –0.40), discomfort due to temperature (cold: r = –0.44; hot: r = –0.39), prolonged sleep latency (r = –0.33), pain (r = –0.25), poor subjective sleep quality (r = –0.24), and nighttime awakenings (r = –0.23). These findings underscore that both the quantity and quality of sleep significantly contribute to the observed multivariate associations. Furthermore, to examine the influence of sleep timing, we applied cosine and sine transformations to bedtime and wake time. The canonical correlation coefficients for the bedtime cosine and sine components were 0.34 and –0.06, respectively, indicating that a bedtime of approximately 23:18 was most strongly associated with the canonical variate. For wake time, the cosine and sine components showed correlations of –0.34 and 0.23, respectively, corresponding to an optimal wake time of approximately 08:50 (Figure 1D).

For cognitive functions, the canonical variate showed strong correlations with cognitive performance measures, particularly overall crystallized cognition (r = 0.79), language (Picture Vocabulary: r = 0.74; Oral Reading Recognition: r = 0.71), fluid intelligence (Penn Progressive Matrices, correct responses: r = 0.71; skipped items: r = −0.69), self-regulation/impulsivity (Delay Discounting, area under the curve for discounting $40,000: r = 0.56, $200: r = 0.53), and spatial orientation (Variable Short Penn Line Orientation correct responses: r = 0.51; total positions off-target: r = −0.56).

The most pronounced anatomical finding was a strong positive association between the canonical variate and volumes of the anterior hippocampus. This was evident in both hemispheres, with the largest correlations observed for the bilateral whole hippocampal head (right: r = 0.77, left: r = 0.88) and molecular layer of the head (right: r = 0.75, left: r = 0.86), and left CA1 head (r = 0.84) (Figure 2B). In contrast, subfields within the hippocampal body and tail generally exhibited weaker, though still positive, or slightly negative correlations. Notably, while initial analyses revealed positive associations between the canonical variate and nearly all hippocampal subfields (Figure 2B), these associations became anatomically specific after controlling for ICV. Specifically, the correlation patterns for sleep and cognition remained unchanged (Figures S1A and S1B), while sustained positive correlations persisted in anterior subfields, and negative correlations emerged in posterior subfields (Figure S1C).

To quantify these anatomical patterns, we regressed the subfield loadings onto their spatial coordinates (Figure 3). This analysis confirmed a strong, significant anterior-posterior gradient; loadings became progressively more positive in more anterior subfields (β = 0.66, p < 0.001; Figure 3A). After accounting for this primary anterior-posterior effect, we also observed a weaker but significant medial-lateral gradient. Specifically, subfields located more laterally tended to have slightly higher loadings than those located more medially (β = −0.25, p = 0.02; Figure 3B).

**Figure 3.**
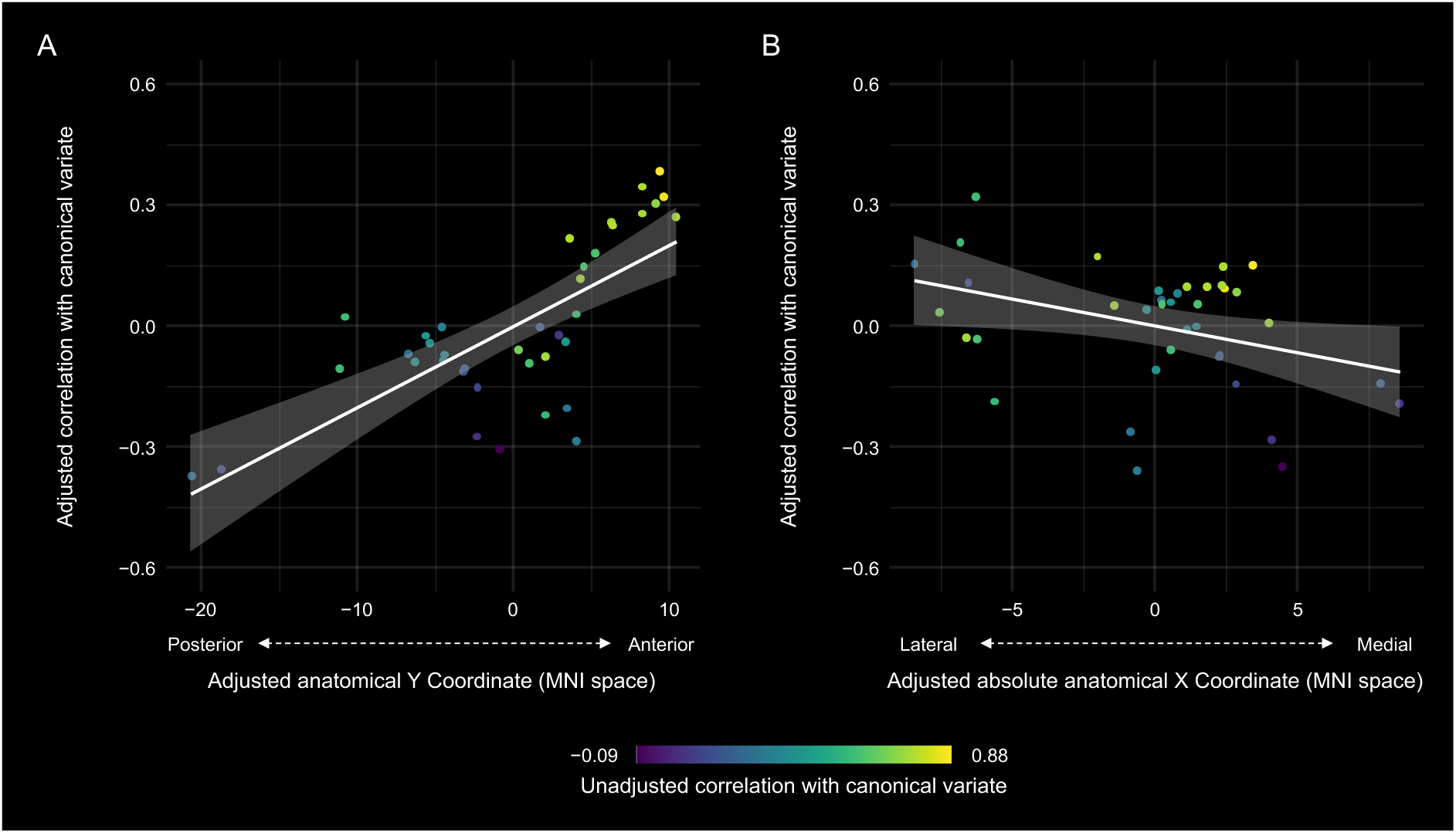
Relationship between hippocampal subfield loadings and their anatomical coordinates. (A) A scatter plot showing a significant positive relationship between the subfield loadings (correlation with the canonical variate) and their position along the anterior-posterior axis (MNI Y-coordinate) after controlling for the hemispheric and medial-lateral effect. (B) A scatter plot showing a significant but weaker relationship between the subfield loadings and their position along the medial-lateral axis (MNI X-coordinate) after controlling for the hemispheric and anterior-posterior effect. In both panels, the white line represents the linear fit, and the shaded area indicates the 95% confidence interval. Each point represents a hippocampal subfield, colored by its original (unadjusted) loading value.

### Sensitivity analysis

All sensitivity analyses confirmed the robustness of our primary findings. We assessed this by comparing the pattern of variable loadings (i.e., the correlations between each original variable and the canonical variate) between our primary model and the various sensitivity models. The correlations with the canonical variate from the primary analysis showed a near-perfect correlation with those from each sensitivity analysis (Pearson’s r = 0.93–0.99). This indicates that the identified mode of covariation and the relative contributions of individual variables are highly stable and not contingent on specific analytical choices. Further, the significant association along the anterior-posterior axis was consistently replicated across all analytical variations, underscoring its robustness. In contrast, the finding for the medial-lateral axis was less consistent, as it failed to reach statistical significance in some sensitivity models.

## Discussion

In this study, we employed a multivariate approach to uncover the complex interplay among sleep, hippocampal subfield anatomy, and cognition in a large cohort of healthy young adults. Our analysis revealed a single, robust mode of covariation: better sleep health—encompassing moderate duration, high quality, and optimal timing—was strongly associated with larger volumes in the hippocampus, an effect most prominent in its anterior aspect. This sleep-hippocampal profile, in turn, was linked to superior performance in higher-order cognitive domains, including fluid intelligence and impulse control. These findings offer compelling evidence that the functional triad of sleep, hippocampal integrity, and cognition is well-established in early adulthood, and highlight an anterior-dominant anatomical pattern in this relationship.

Consistent with a large body of previous research, our results underscore the importance of multiple sleep dimensions for brain health. We identified a nonlinear, inverted U-shaped association between sleep duration and the canonical variate, which aligns with and extends previous findings that linked both very short and long sleep to smaller total hippocampal volumes (Kuperczkó et al., 2015; Tai et al., 2022). This nonlinear relationship suggests that a moderate sleep duration is most beneficial for the integrated profile of hippocampal structure and cognitive function. Physiologically, this optimal range may reflect a balance essential for synaptic homeostasis, where sufficient sleep is required for synaptic downscaling and memory consolidation, while excessive sleep may indicate underlying health issues or fragmented, poor-quality rest (Grandner & Drummond, 2007; Tononi & Cirelli, 2014). Moreover, our results emphasize that it is not merely the quantity but also the restorative quality of sleep that is critical for brain health, as metrics like fewer awakenings and less discomfort were key contributors to the healthy sleep profile. Our findings on sleep timing further reinforce the principle of circadian alignment. The optimal bedtime we identified (around 23:18) is slightly later than that in a large Chinese cohort (Ouyang et al., 2024) but earlier than the late bedtimes associated with smaller hippocampal volumes in Hungarian young adults (Kuperczkó et al., 2015). While these discrepancies may reflect cultural or demographic differences (Ou et al., 2025), they converge on the idea that adherence to a regular, appropriately timed sleep schedule is crucial for supporting the molecular and cellular processes that underpin hippocampal plasticity and resilience.

Crucially, our study moves beyond whole-hippocampus analyses by revealing that these sleep-related associations are not uniform across the hippocampus, but rather show a pronounced anterior-posterior gradient. The hippocampus is not a uniform structure; it exhibits significant functional specialization along its longitudinal axis (Strange et al., 2014). The posterior hippocampus is primarily implicated in fine-grained spatial navigation and perceptual details, whereas the anterior hippocampus, the region most strongly implicated in our study, is considered a hub for processing emotion, stress, motivation, and more flexible, gist-based memory representations (Poppenk et al., 2013; Strange et al., 2014). It has dense connections with the amygdala and prefrontal cortex, which allows it to integrate affective and cognitive information. The pronounced association between sleep and the anterior hippocampus in our results is therefore highly significant. This anatomical gradient suggests that healthy sleep may be especially important for supporting anterior hippocampal functions in young adults. For instance, many of the sleep quality metrics that loaded strongly in our model (e.g., discomfort, pain) are related to stress and affective state, which are primarily processed by the anterior hippocampus and its connected circuits. Furthermore, the cognitive functions most strongly associated with the sleep-hippocampal signature—language, fluid intelligence, self-regulation/impulsivity, and spatial orientation—are precisely the types of higher-order, flexible cognitive processes attributed to the anterior hippocampus and its interactions with the prefrontal cortex (Rubin et al., 2014). Thus, our findings suggest a consistent neuroanatomical pattern, linking the restorative aspects of sleep to the structure of a key cognitive-affective hub, which in turn supports complex cognitive abilities.

The significance of this anatomical gradient is further underscored by our analysis controlling for ICV. Better sleep has been associated with greater brain volume more broadly (Tai et al., 2022). The disappearance of the positive association with the posterior hippocampus after controlling for ICV likely reflects the removal of this general, non-specific effect of sleep on brain morphology. Notably, the persistence of a positive association with the anterior hippocampus, even after accounting for ICV, may suggest a regionally specific relationship between sleep and anterior hippocampal structure. This anterior-dominant pattern raises the possibility that sleep could be particularly relevant for supporting functions subserved by the anterior hippocampus.

Our integrated findings illuminate how sleep quality, specific hippocampal subfield structures, and higher-order cognitive functions are interrelated in healthy young adults, addressing a previously underexplored area in the literature. While previous studies have linked sleep to hippocampal subfields in clinical or aging populations (Joo et al., 2014; Liu et al., 2021), our work demonstrates that this relationship is present and functionally significant in a large, well-characterized, non-clinical sample of young adults. Our finding of anterior hippocampal–cognitive covariation reinforces the idea that this region is a vital substrate for sleep-dependent cognitive processes throughout the lifespan (Teicher et al., 2017). It suggests that the foundations of cognitive resilience in later life may be partly established through sleep-related hippocampal maintenance in early adulthood.

Our study has several strengths, including the large, well-characterized HCP sample, the use of high-resolution subfield segmentation, and the application of MCCA, which allowed us to model the complex, multivariate nature of the sleep-brain-cognition nexus. However, some limitations must be acknowledged. First, the cross-sectional design precludes any causal inferences. Longitudinal studies are needed to determine whether poor sleep leads to hippocampal changes, vice versa, or if a third factor influences both. Second, our reliance on the self-reported PSQI, while widely used, is subject to reporting bias. Future studies incorporating objective sleep measures, such as actigraphy or polysomnography, would provide a more complete picture. Finally, our healthy, young adult sample limits generalizability to other age groups or clinical populations.

## Conclusion

This study provides a nuanced and integrated view of the relationship between sleep, hippocampal structure, and cognition. By demonstrating a robust association between better sleep, larger anterior hippocampal volumes, and enhanced cognitive function in healthy young adults, we highlight the importance of sleep for brain health early in the lifespan. Our findings highlight an anterior-posterior gradient of sensitivity within the hippocampus, pinpointing the anterior subfields as a key modulator of the link between sleep and cognition. These findings challenge the view that sleep-related brain changes are confined to later life and provide a specific neuroanatomical target for future research and potential interventions aimed at preserving cognitive function across the lifespan by promoting healthy sleep.

## Acknowledgments

Data were provided by the Human Connectome Project, WU-Minn Consortium (Principal Investigators: David Van Essen and Kamil Ugurbil; 1U54MH091657) funded by the 16 NIH Institutes and Centers that support the NIH Blueprint for Neuroscience Research; and by the McDonnell Center for Systems Neuroscience at Washington University.

## Authors’ contributions

Tianfang Han: Conceptualization, Data Curation, Formal Analysis, Visualization, Writing – Original Draft, Funding Acquisition. Tongfang Ding: Data Curation, Writing – Review & Editing. Shubing Li: Writing – Review & Editing. Toru Ishihara: Conceptualization, Methodology, Formal Analysis, Investigation, Resources, Supervision, Project Administration, Writing – Review & Editing, Funding Acquisition.

## Data availability statement

All data used in this study are publicly available from the Human Connectome Project database (https://www.humanconnectome.org/).

---

**Supplementary Table 1.** Description of cognitive tests and variables.

| Test | Variable Index | Cognitive Domain | Outcome Measurement Description |
| --- | --- | --- | --- |
| Picture Sequence Memory Test | PicSeq_Unadj | Episodic Memory | Number of correctly recalled adjacent picture pairs |
| Dimensional Change Card Sort Test | CardSort_Unadj | Executive Function / Cognitive Flexibility | Score based on accuracy and reaction time |
| Flanker Inhibitory Control and Attention Test | Flanker_Unadj | Executive Function / Inhibitory Control | Score based on accuracy and reaction time |
| Penn Matrix Test | PMAT24_A_CR | Fluid Intelligence | Number of correct responses |
|  | PMAT24_A_SI | Fluid Intelligence | Number of skipped items |
|  | PMAT24_A_RTCR | Fluid Intelligence | Median reaction time for correct responses |
| Oral Reading Recognition Test | ReadEng_Unadj | Language / Reading Decoding | Score based on reading accuracy |
| Picture Vocabulary Test | PicVocab_Unadj | Language / Vocabulary Comprehension | Score from a computer-adaptive vocabulary test |
| Pattern Comparison Processing Speed Test | ProcSpeed_Unadj | Processing Speed | Number of correct responses |
| Delay Discounting | DDisc_AUC_200 | Self-regulation / Impulsivity | Area under the curve for a delayed reward of \$200 |
| | DDisc_AUC_40K | Self-regulation / Impulsivity | Area under the curve for a delayed reward of \$40,000 |
| Variable Short Penn Line Orientation Test | VSLOT_TC | Spatial Orientation | Total number of correct responses |
|  | VSLOT_CRTE | Spatial Orientation | Median reaction time on correct trials, adjusted for task difficulty |
|  | VSLOT_OFF | Spatial Orientation | Total positions off-target across all trials |
| Short Penn Continuous Performance Test | SCPT_TP | Sustained Attention | Number of true positives |
|  | SCPT_TN | Sustained Attention | Number of true negatives |
|  | SCPT_FP | Sustained Attention | Number of false positives |
|  | SCPT_FN | Sustained Attention | Number of false negatives |
|  | SCPT_TPRT | Sustained Attention | Median reaction time for true positive responses |
|  | SCPT_SEN | Sustained Attention | Sensitivity (true positive rate) |
|  | SCPT_SPEC | Sustained Attention | Specificity (true negative rate) |
|  | SCPT_LRNR | Sustained Attention | Longest run of non-responses (measure of lapses) |
| Penn Word Memory Test | IWRD_TOT | Verbal Episodic Memory | Total number of correctly recalled words |
|  | IWRD_RTC | Verbal Episodic Memory | Median reaction time for correct recalls |
| Emotion Recognition Test | ER40_CR | Emotion Recognition | Total number of correct responses |
|  | ER40_CRT | Emotion Recognition | Median reaction time for correct responses |
|  | ER40ANG | Emotion Recognition | Number of correct identifications of "angry" |
|  | ER40FEAR | Emotion Recognition | Number of correct identifications of "fear" |
|  | ER40HAP | Emotion Recognition | Number of correct identifications of "happy" |
|  | ER40NOE | Emotion Recognition | Number of correct identifications of "neutral" |
|  | ER40SAD | Emotion Recognition | Number of correct identifications of "sad" |
| List Sorting Working Memory Test | ListSort_Unadj | Working Memory | Number of correctly sorted lists |
| Composite Scores | CogFluidComp_Unadj | Fluid Cognition | Composite score of fluid cognitive abilities |
|  | CogEarlyComp_Unadj | Early Childhood Cognition | Composite score of early childhood cognitive abilities |
|  | CogTotalComp_Unadj | Overall Cognition | Composite score of overall cognitive function |
|  | CogCrystalComp_Unadj | Crystallized Cognition | Composite score of crystallized cognitive abilities |

**Figure S1.**
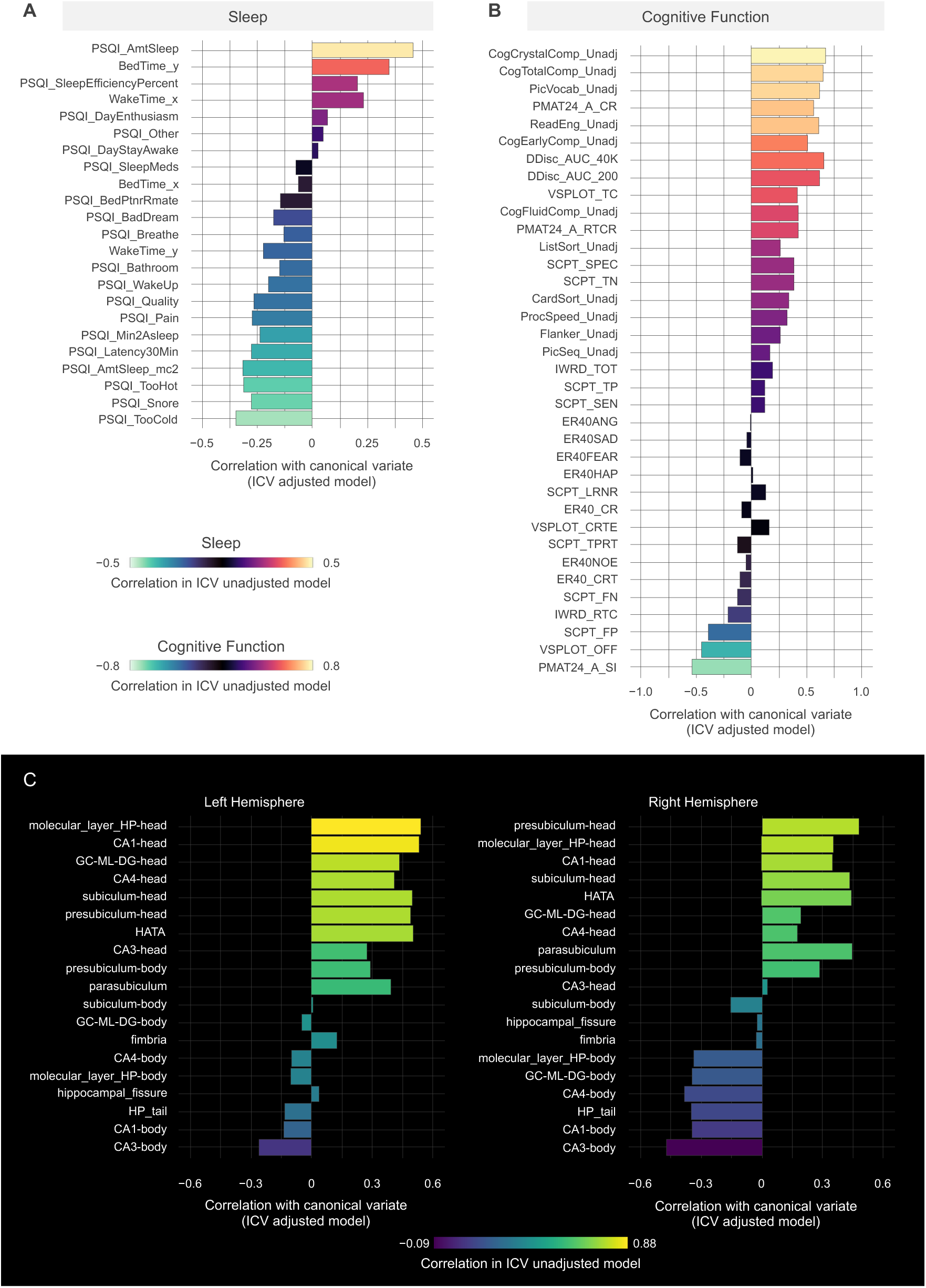
Mode of covariation among sleep, cognitive function, and hippocampal volume after controlling for intracranial volume (ICV). To facilitate comparison with the primary analysis, the bars in all panels are ordered and colored according to their corresponding loadings in the unadjusted model (as shown in Figures 1 and 2). (A, B) Correlations of individual sleep (A) and cognitive function (B) parameters with their respective canonical variates. The loading patterns are nearly identical to the primary analysis (Figure 1). (C) Correlations of individual hippocampal subfield volumes with the hippocampal canonical variate. After controlling for ICV, the association becomes anatomically specific: positive correlations remain in anterior subfields, while negative correlations emerge in posterior subfields.

